# The balance of acidic and hydrophobic residues predicts acidic transcriptional activation domains from protein sequence

**DOI:** 10.1101/2023.02.10.528081

**Authors:** Sanjana R. Kotha, Max Valentín Staller

## Abstract

Transcription factors activate gene expression in development, homeostasis, and stress with DNA binding domains and activation domains. Although there exist excellent computational models for predicting DNA binding domains from protein sequence (Stormo, 2013), models for predicting activation domains from protein sequence have lagged behind (Erijman et al., 2020; Ravarani et al., 2018; Sanborn et al., 2021), particularly in metazoans. We recently developed a simple and accurate predictor of acidic activation domains on human transcription factors (Staller et al., 2022). Here, we show how the accuracy of this human predictor arises from the balance between hydrophobic and acidic residues, which together are necessary for acidic activation domain function. When we combine our predictor with the predictions of neural network models trained in yeast, the intersection is more predictive than individual models, emphasizing that each approach carries orthogonal information. We synthesize these findings into a new set of activation domain predictions on human transcription factors.

## Introduction

Transcription factors regulate gene expression with DNA binding domains and effector domains. DNA binding domains are structured, conserved, and recognize related DNA sequences (Ferrie et al., 2022; Latchman, 2008). Profile hidden Markov models can accurately predict DNA binding domains from protein sequence (El-Gebali et al., 2019; Finn et al., 2016; Stormo, 2013). Effector domains include repression domains that bind corepressors and activation domains that bind coactivators. Some repression domains can be predicted from protein sequence, and many contain short linear motifs (DelRosso et al., 2022; Soto et al., 2022; Tycko et al., 2020). Activation domains are intrinsically disordered, poorly conserved, and bind to structurally diverse coactivators: these features have made it difficult to predict activation domains from protein sequence. There are profile hidden Markov models for individual activation domains (e.g. p53 or Hif1a), which can identify activation domains on closely related paralogs or orthologs in other vertebrate species (El-Gebali et al., 2019), but these models are rarely generalizable to predict activation domains on other transcription factors. The ability to predict activation domains from protein sequence would open the door to automated annotation of proteomes and lay a foundation for prioritizing disease-causing mutations. Recently developed models trained in yeast have provided a useful starting point (Erijman et al., 2020; Ravarani et al., 2018; Sanborn et al., 2021). Here, we explore how to improve these models to work on human transcription factors.

Over the last few years, we and others have resolved the key sequence features of strong acidic activation domains, the largest known class (Arnold et al., 2018; Broyles et al., 2021; DelRosso et al., 2022; Erijman et al., 2020; Ravarani et al., 2018; Sanborn et al., 2021; Staller et al., 2022, 2018; Tycko et al., 2020). Based on these sequence features, we proposed an *acidic exposure model* for activation domain function (Staller et al., 2022, 2018). In our acidic exposure model, the hydrophobic residues make key contacts with coactivators, but left alone, the hydrophobic residues interact with each other and prevent binding to partners (**Figure 1A**). Interspersed between hydrophobic residues are acidic residues that repel each other and keep the hydrophobic residues exposed to solvent. The critical parameter in the acidic exposure model is the balance between acidic and hydrophobic residues. There are cases where acidic residues make fast, low-affinity contacts with basic residues on coactivators (Almlof et al., 1997; Hermann et al., 2001; Kim and Chung, 2020), but these are secondary to hydrophobic contacts.

**Figure 1:**
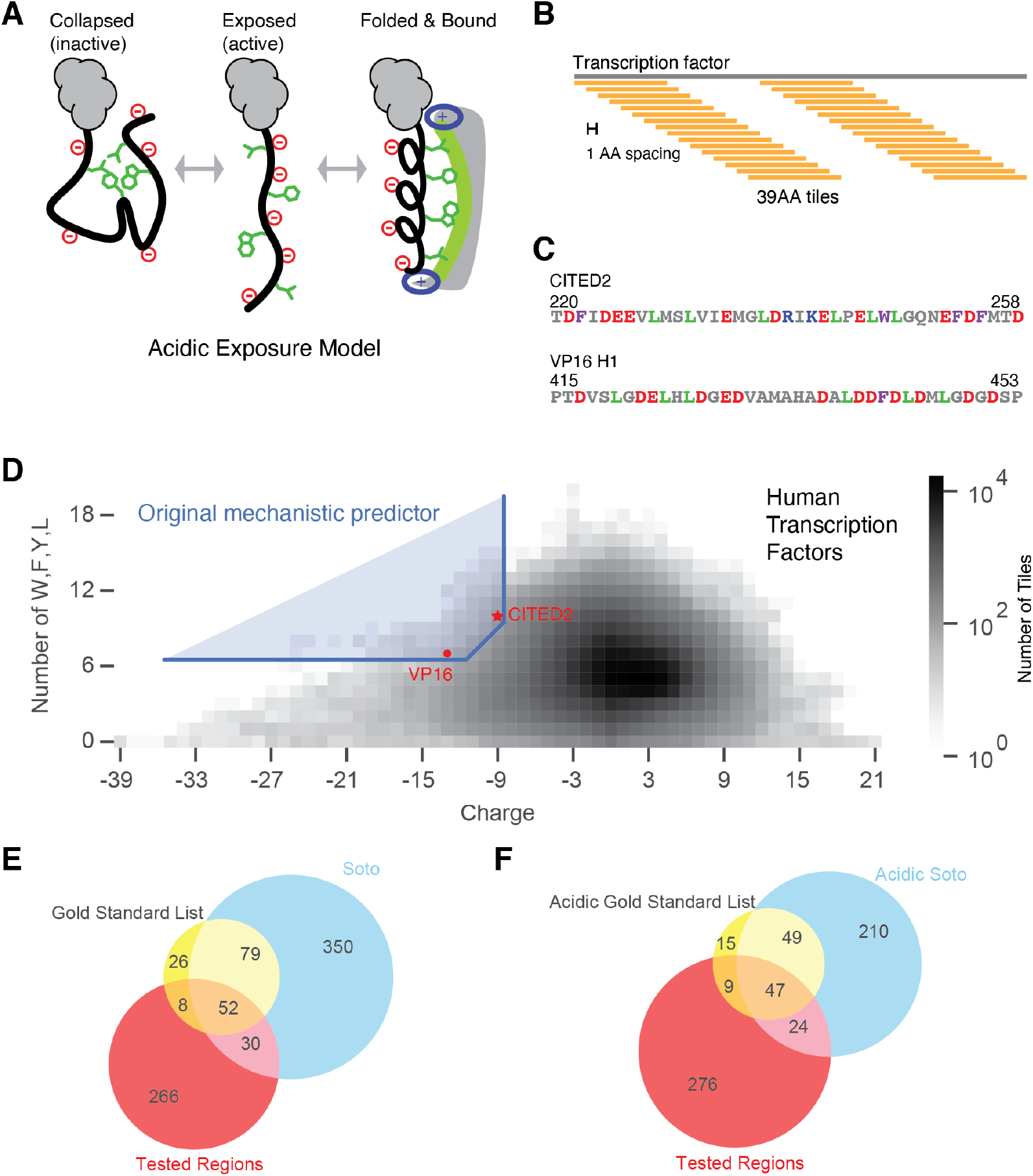
The acidic exposure model can predict acidic activation domains on human transcription factors. A) In the acidic exposure model, intrinsically disordered activation domains dynamically morph between a collapsed, inactive state and an expanded, active state where the key aromatic and leucine residues are available to interact with hydrophobic surfaces of coactivators. Many activation domains show coupled folding and binding. B) Schematic for how we took transcription factors and computationally chopped them into 39-AA tiles spaced every 1-AA. C) Sequences of a 39-AA version of CITED2 and a 39-AA version of VP16 H1 used in the predictor. D) The original mechanistic predictor, blue boundary, is anchored by CITED2 (red star) and VP16 H1 (red circle). E) Overlap between the gold standard list, the Soto list, and the Tested Regions from Staller et al., 2022. The tested regions include predictions, known activation domains, and negative control regions. F) Overlap between the acidic subsets of the gold standard list, the Soto list, and tested regions.

The acidic exposure model motivated a simple, interpretable, and accurate mechanistic predictor of activation domains (Staller et al., 2022). Our rational mutagenesis of CITED2 (220-258) and VP16 H1 (415-453) revealed that the key sequence features controlling activity of these activation domains were net negative change and the number of aromatic and leucine residues. Both these proteins lack a DNA binding domain: VP16 H1 is from the human herpes simplex virus while CITED2 is often labeled a coactivator (Freedman et al., 2003; Sadowski et al., 1988). We found that regions that resemble CITED2 and VP16 H1 in acidity and the number of aromatic and leucine residues were enriched for known and new activation domains (Staller et al., 2022). First, we computationally decomposed 1608 human transcription factors into 39 amino acid (39-AA) tiles spaced every 1-AA, yielding 881K tiles (**Figure 1B**) (Lambert et al., 2018). We looked for 39-AA tiles that were similar to VP16 H1 and CITED2 (**Figure 1C)**, using the the formula:

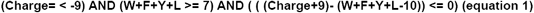

Where W is tryptophan, F is phenylalanine, Y is tyrosine, and L is leucine. This predictor is anchored by CITED2 (net charge = -9, W+F+Y+L = 10) and VP16 H1 (net charge = -13, W+F+Y+L = 7) (**Figure 1D**). The first term selected regions that are at least as acidic as CITED2 (**Figure 1D**, vertical threshold). The second term selected regions with at least as many W,F,Y, or L residues as VP16 H1 (**Figure 1D**, horizontal threshold). The third term interpolates between these two anchor points with a diagonal line of slope 1 (**Figure 1D**, diagonal line). In the human proteome, 1139 39-AA tiles met these criteria, which combined into 144 predicted activation domains. Twenty-six of these predictions overlapped known activation domains, more than expected by chance (p < 1e-5 in permutation tests). We split the longest predictions and tested 150 regions in our high-throughput assay (Staller et al., 2022). Of these, 108/149 (72%) had detectable activity and 58/149 (39%) had high activity. This fraction compared favorably to length-matched random regions and positive controls. These hits included 28 known activation domains and 30 new activation domains (Staller et al., 2022).

While we were developing our mechanistic predictor for human activation domains, two convolutional neural networks for predicting activation domains in yeast were published. The first, ADpred, was trained on 3.6 million 30-AA random peptides (Erijman et al., 2020). The second, PADDLE, was trained on 53-AA regions that tiled across ∼180 *Saccharomyces cerevisiae* transcription factors (n = 7460 tiles) (Sanborn et al., 2021). Both of these datasets see the same primary signal we saw in our rational mutagenesis: strong activation domains are enriched for acidic and aromatic residues and depleted of basic residues. In yeast, four groups have reported the same ranking of amino acid contributions to activity: W > F > Y > L (Erijman et al., 2020; Ravarani et al., 2018; Sanborn et al., 2021; Staller et al., 2018). Sanborn et al. reported a correlation between White-Wimbly hydrophobicity and activity, but in this hydrophobicity table the top entries are W > F > Y > L, so the White-Wimbly hydrophobicity scale is emphasizing the most important amino acids (Sanborn et al., 2021). Erijman et al. found that [D,E][W,F,Y] dipeptides were enriched in active fragments (Erijman et al., 2020). Acidic residues collectively contribute to activity by creating a permissive context. Both of these signals support the acidic exposure model and are consistent with our mechanistic predictor.

Here, we sought to understand why our mechanistic predictor accurately predicted transcriptional activation domains on human transcription factors and to improve its predictive power. Using a new, hand-curated gold standard list, we found that the key feature that provides predictive power is the balance between acidic residues and specific hydrophobic residues. Phenylalanine and leucine made the largest contributions to predictive power, while tryptophan, tyrosine, and methionine contributed modestly. Our new predictor uses tryptophan, phenylalanine, and leucine residues, 39-AA windows, and more relaxed thresholds for net charge and hydrophobic residue counts. When we compare our original mechanistic predictor to convolutional neural network models trained in yeast, we found that the intersection is more predictive than individual models, emphasizing that each approach carries distinct information. We synthesize these findings into a new set of activation domain predictions.

## Results

To quantify the performance of our activation domain predictors, we curated a gold standard list of 167 activation domains from 135 proteins, 129 on human transcription factors (**Table S1**). This list combined Uniprot annotated activation domains (Downloaded June 2020), individual activation domains curated from NMR papers, and a classic hand-curated list of activation domains (Choi et al., 2000). Overlapping entries were combined by taking the lower start and greater end to make longer annotations. Activation domain boundaries remain difficult to define, so we chose permissive boundaries. When looking for overlaps betweens lists of activation domains, we started with a very permissive threshold, ≥1 overlapping residue, but the minimum observed overlap was 26 residues. We developed our predictors using this gold standard list and validated the predictors with another recently published “Soto list” (Soto et al., 2022). To avoid circular reasoning, the validation set did not include the 30 novel activation domains correctly identified by the original predictor (Staller et al., 2022). These 30 continue to be correctly identified by our modified predictors. Our gold standard list and the Soto list are highly overlapping (**Figure 1E**), and neither list represents a complete list of true positives.

Documented activation domains are more diverse than the traditional categories of acidic, proline-rich, or glutamine-rich (Gerber et al., 1994; Latchman, 2008; Sigler, 1988). There have been scattered references to alanine-rich, glycine-rich, and serine-rich activation domains in the literature, but they have not been recognized as archetypes (Alerasool et al., 2022; Schaeffer et al., 1999; Soto et al., 2022). Using a threshold of 15% for composition bias, our gold standard list contains 105 (62.9%) acidic (net charge < -3), 12 (7.19%) glutamine-rich (Q-rich), and 30 (18.0%) proline-rich (P-rich) activation domains. In addition, there are 37 (22.2%) serine-rich (S-rich) and 7 alanine-rich (4.19%) (**Table 1**). The Soto et al. list contains 290 (79.6%) acidic, 18 (3.5%) Q-rich, 119 (22.9%) P-rich, 124 (23.8%) S-rich, 36 (6.9%) glycine-rich, and 9 (1.7%) alanine-rich activation domains. Using our 15% criteria, some activation domains are enriched for more than one amino acid. Annotated acidic activation domains on the two lists also overlap with our tested regions (**Figure 1F**). Activation domains are enriched for disorder promoting residues, consistent with the evidence that nearly all activation domains are intrinsically disordered (Hahn and Young, 2011; Liu et al., 2006; Oldfield and Dunker, 2014; van der Lee et al., 2014). We confirmed that >90% of the activation domains on both lists are predicted to be intrinsically disordered by Metapredict2 (**Figure S1**) (Emenecker et al., 2022).

**Table 1:** Types of activation domains on the Gold Standard List (GSL) and Soto et al. List.

|  | Number of GSL Entries | Number of Soto Entries |
| --- | --- | --- |
| Acidic | 105 | 290 |
| Glutamine-rich | 12 | 18 |
| Proline-rich | 30 | 119 |
| Serine-rich | 37 | 124 |
| Alanine-rich | 7 | 9 |

We examined the sequence features of our gold standard list of activation domains. As a background distribution, we used a published list of 1608 human transcription factors (Lambert et al., 2018). Annotated activation domains have a wide distribution of lengths because only some have been experimentally minimized (**Figure 2**). To make our analyses more consistent, we performed all composition analysis by decomposing each activation domain into all possible 39-AA sliding windows, spaced at 1-AA intervals (e.g. a 45 residue activation domain region would become 7 39-AA tiles, **Figure 1B**). We started with 39-AA tiles because that was the length-scale of the original predictor. These 39-AA tiles accommodate activation domains of different lengths and avoid the difficult problem of defining activation domain boundaries. Many activation domains are 39AA or shorter and many long ones contain highly active subregions. Throughout this work we will analyze proteins by decomposing them into 39-AA tiles and comparing the features of these sets of tiles. Compared to human transcription factor tiles, activation domain tiles show a modest enrichment of net negative charge and M, D, S, L, P, Q, Y, A, V, G residues (t-test, p < 1e-4, Bonferroni corrected, **Figure 2, Figure S2**). Activation domains from the gold standard list do not exhibit extreme properties compared to the background sequence properties of human transcription factors (**Figure S2, S3**). This similarity to the background distribution explains in part why activation domains have been so difficult to predict from sequence.

**Figure 2:**
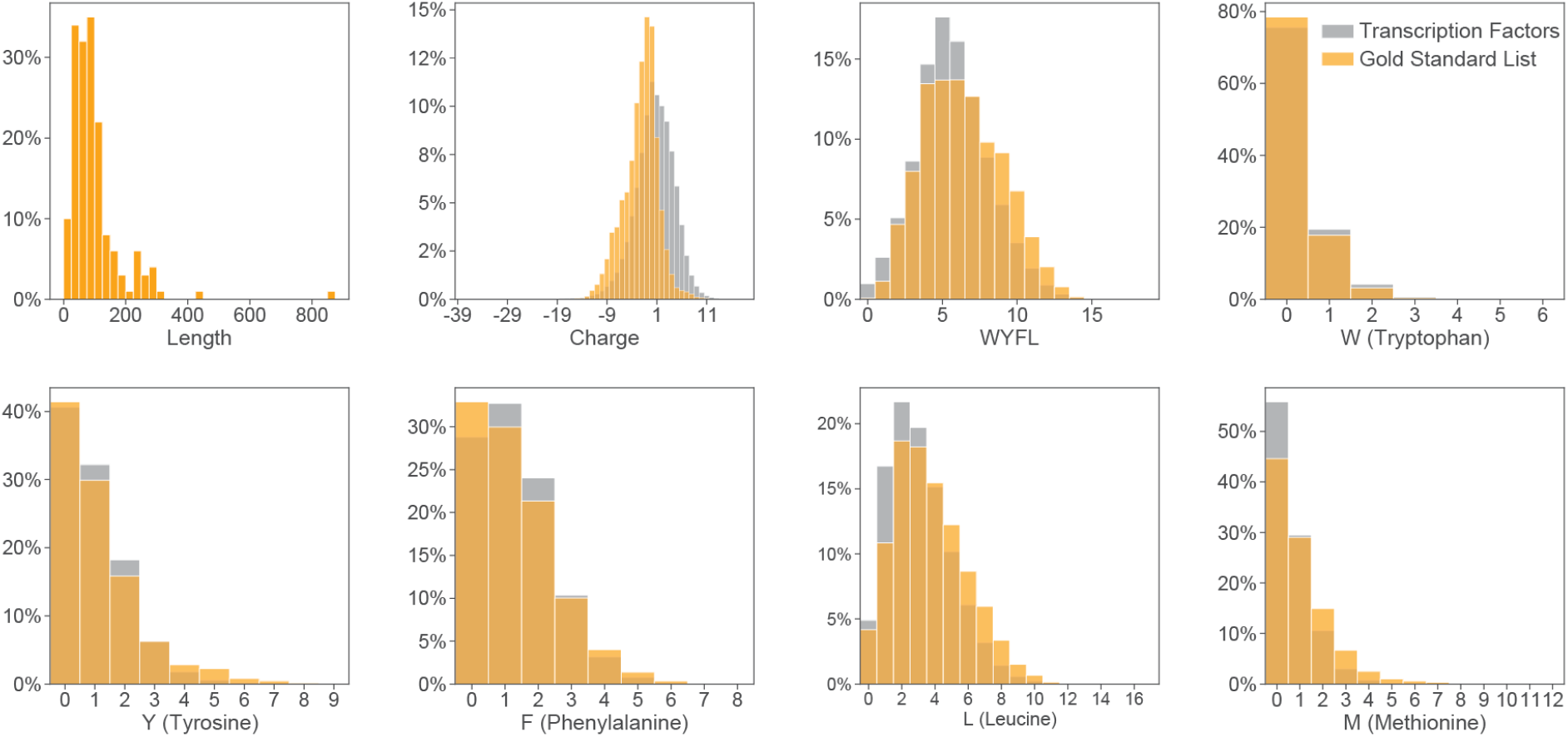
Sequence features of activation domains. Activation domains on the gold standard list (orange) show very moderate enrichment of individual sequence properties or amino acids compared to the background of transcription factor sequences (gray).

We next examined how our mechanistic predictor accurately identified acidic activation domains using the combination of net charge and the number of W+F+Y+L residues. To establish a background distribution, we first examined the sequence features of the full human proteome (**Figure 3A**) and 1608 human transcription factors (Lambert et al., 2018), **Figure 3B**). Compared to the full human proteome, transcription factor tiles are slightly positively charged, likely because DNA binding domains often contain many basic residues that electrostatically interact with the acidic phosphate backbone (**Figure 3D**). Transcription factors contain fewer W+F+Y+L residues than the full proteome, likely because they are depleted for transmembrane domains and globular folded protein cores (**Figure 3E**). Transcription factors contain strong local biases in net charge (**Figure 3B**). The most common tile net change for transcription factors and the proteome is neutral (**Figure 3E**). The distribution of transcription factor tile properties is reasonably representative of the full proteome.

**Figure 3:**
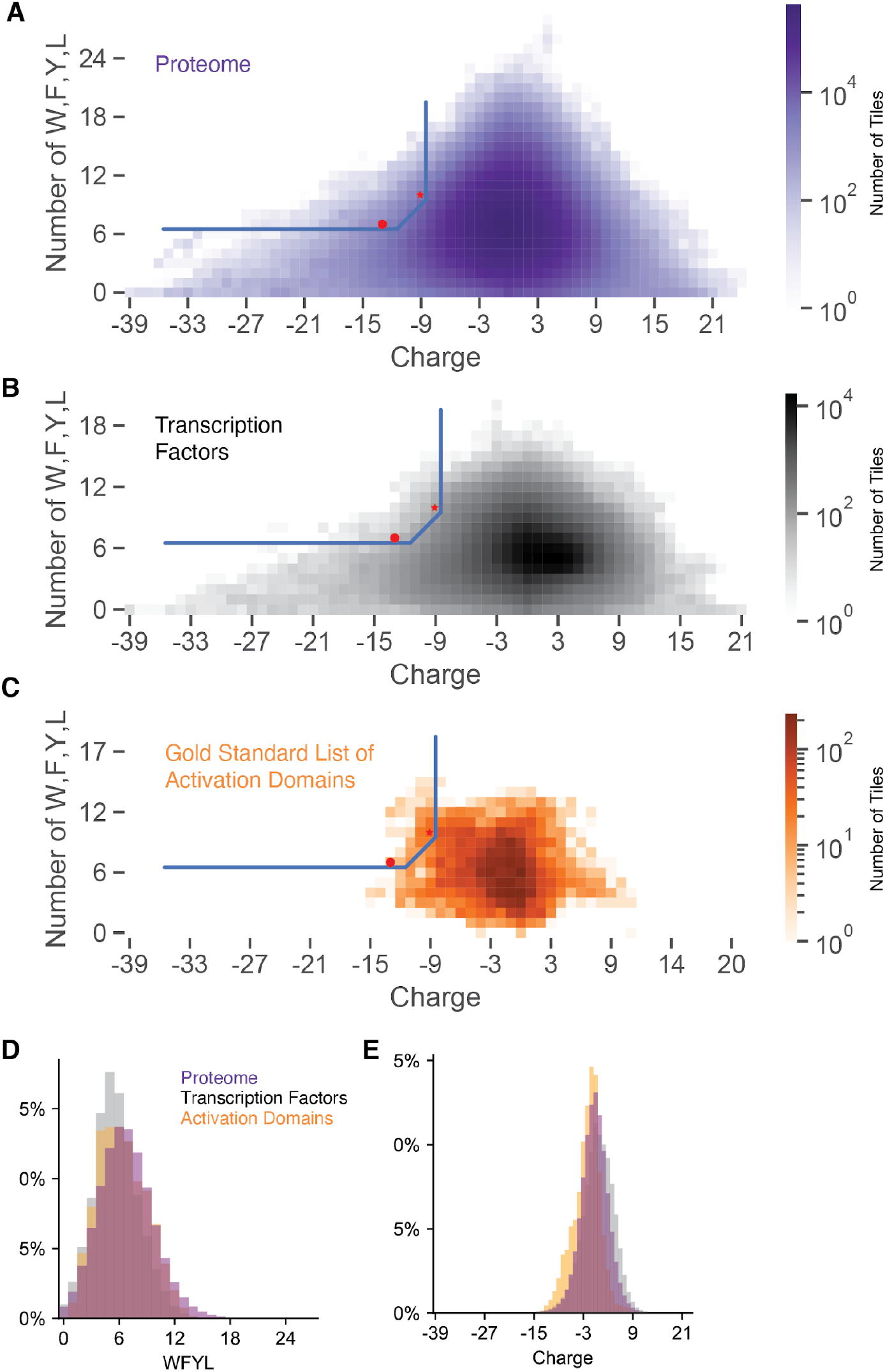
The combination of acidic and WFYL residues predicts activation domains. A) We decomposed the proteome into 39AA tiles and calculated the net charge and counted WFYL residues for each tile. For each combination of these two properties, we counted tiles to create a 2D histogram that is visualized as a heatmap. The full proteome has a more diverse distribution of tiles than other protein sets we examined. B) Tiles from annotated transcription factors. C) Tile from the gold standard list of activation domains. D) Overlaps of histograms of tiles with varying numbers of WFYL residues. E) The gold standard list of activation domains is enriched for acidic tiles.

Long runs of acidic amino acids are far more common than runs of basic amino acids (Bigman et al., 2022). There exist 11 tiles from transcription factors with a charge of -39 (D/E runs), spanning residues 258-307 of MYT1, a neural transcription factor (Nielsen et al., 2004). Conversely, the six most positively charged tiles from transcription factors were +21, which spanned residues 1845 - 1908 of SON, a splicing cofactor that binds DNA (Mattioni et al., 1992). The net charge of proteome tiles is also asymmetric, spanning -39 to +24. The most acidic patch in the proteome is 50 consecutive acidic residues (-50), but the most basic region is +24 (Bigman et al., 2022).

When we compared the sequence properties of tiles of activation domains to tiles of full length transcription factors, we found that acidic activation domains are more acidic than transcription factors (**Figure 3E**) and have a slight enrichment for W+F+Y+L residues (**Figure 3D**). However, the enrichment for the combination of acidity and W+F+Y+L residues is much stronger (**Figure 3C**). The most acidic regions of transcription factors are not part of known activation domains. Although the most acidic regions are visible on a log scale (**Figure 3C**), they are rare and not visible on a linear scale (**Figure 3E**). Similarly, the transcription factor tiles with the most W+F+Y+L residues are not part of activation domains (**Figure 3C, 3D**). No single property distinguishes activation domains, but the combination of acidity and W+F+Y+L residues can enrich a subclass of acidic activation domains. Our predictor finds activation domains that balance acidic residues against W+F+Y+L residues (**Figure 3C**).

Similarly to how acidic activation domains are not the most acidic regions of transcription factors, P-rich and Q-rich activation domains are not among the transcription factor tiles with the most P’s or Q’s (**Figure S4, S5**). This observation is consistent with evidence that Q-rich activation domains contain a lower fraction of Q’s than proteins with true poly-Q regions, like Huntingtin (Ruff et al., 2014). The traditional activation domain labels were assigned before completion of the human genome project, namely before there was a proper null distribution against which to show enrichment.

In contrast, S-rich activation domains are enriched for serine residues compared to the background of transcription factor sequences. The tiles spanning S-rich activation domains on our gold standard list had more serine residues than tiles from all other regions of transcription factors (t-test, pval = 1.37e-229). Some activation domains increase activity when they are phosphorylated (Conti et al., 2023; De Mol et al., 2018; Peng et al., 2019; Raj and Attardi, 2017), prompting us to search for an enrichment of common phosphorylation motifs, (e.g. SP and SQ). Activation domains and repression domains contain more documented phosphorylation sites than DNA binding domains (Soto et al., 2022). We initially detected an enrichment of these motifs when we compared the gold standard list to all transcription factor tiles. As a more stringent control, we shuffled the sequences of the S-rich activation domains and counted spurious occurrences of phosphorylation motifs. Compared to these sequence permutations, phosphorylation motifs did not occur in activation domains more often than expected by chance. We conclude that phosphorylation motifs are not enriched in S-rich human activation domains.

We tested whether the combination of W+F+Y+L and P, Q, or S could enrich activation domains of each class, but none of these combinations worked (**Figure S4, S5, S6**). Our simple, mechanistic predictor of acidic activation domains is not easily extended to other classes.

### Dissecting the mechanistic predictor

In order to improve the predictor, we next sought to understand why it worked and which activation domains from the gold standard list it was identifying. We first asked if human activation domains were more similar to VP16 H1 or the CITED2 by dividing our initial prediction region into three triangular regions (**Figure 4A, Table 2**). Region A has the highest fraction of correct predictions (12/47). Region B, where acidic and hydrophobic residues were balanced, had the most correct predictions (25/133). In contrast, Region C identified no activation domains on the gold standard list (0/18). Tiles like CITED2 and balanced tiles were most likely to be activation domains. This analysis promoted us to remove Region C from the updated predictor.

**Figure 4:**
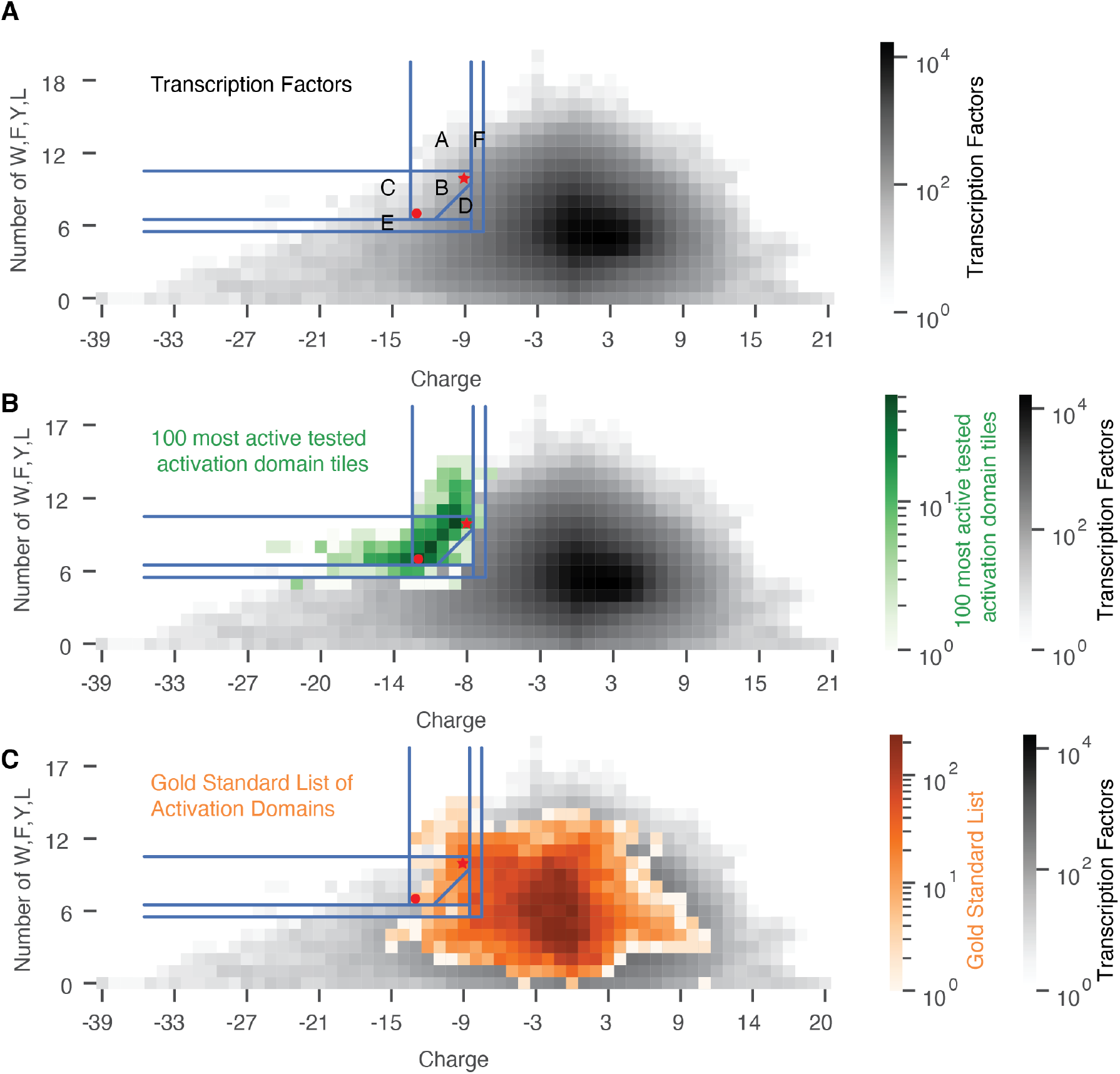
Regions that balance acidity and WFYL residues are most predictive of activation domains. A) We split our original activation domain predictor into regions A, B, and C. We tested the predictive power of additional regions D, E, and F. The red dot indicates VP16 H1, and the red star indicates CITED2, the anchor points for the original activation domain predictor. B) Projecting the tiles from the 100 strongest active activation domains onto regions in A. C) Projecting tiles of the gold standard list over the regions in A.

**Table 2:** Subregions of the mechanistic predictor from Figure 3A differ in the power to detect members of the Gold Standard List (GSL) of activation domains. Source: Table 2 Notebook

| Region | Number of Predictions | GSL Overlap Count | GSL Overlap Proportion |
| --- | --- | --- | --- |
| A | 47 | 12 | 0.255 |
| B | 133 | 25 | 0.188 |
| C | 18 | 0 | 0.000 |
| D | 285 | 28 | 0.982 |
| E | 202 | 7 | 0.035 |
| F | 633 | 50 | 0.079 |

Reciprocally, when we took our correct predictions (i.e. predictions with high activity in our experiment (Staller et al., 2022) and examined their tiles, the peak of distribution lay along the line connecting CITED2 and VP16 (**Figure 4B**). Virtually all of the tiles that mapped to Region C came from activation domains that also contained tiles that mapped to Region B. This analysis further emphasizes how balance is the key to accurate prediction.

To determine the parameters that contribute most to the predictor, we performed sensitivity analysis. We removed each of the eight AAs in the predictor and recomputed predictive power (**Table S2**). F, L, and charge make the largest contributions to sensitivity and specificity, likely because these residues are more common. Despite being enriched in the gold standard list activation domains, Y made very small contributions. W’s make modest contributions to predictive power because they are rare. Similarly, we varied the length of the tiling windows and did not see improvement (**Table S3**). There was no simple way to improve upon the original predictor.

For further comparison, we replaced the y-axis parameter with all singles, pairs, and triplets of amino acids (**Table S4, Table S5, Table S6**). We changed the thresholds based on the sequences of CITED2 and VP16 H1 (methods). Leucine was the single amino acid with the highest specificity. Leucine was present in the five pairs with the highest specificity and in 11/12 triplets with the highest specificity. Together, this analysis emphasized that a high number of leucine residues is predictive of human activation domains.

### Expanding the boundaries of the mechanistic predictor

To make new predictions, we altered the boundaries of the mechanistic predictor to include more tiles. First, we changed the interpolation between VP16 H1 and CITED2 from a diagonal line to a corner, using the minimum value of each activation domain to create a right triangle below CITED2 (**Figure 4A, Region D, Table 3**) using equation 2.

**Table 3:** Changing the interpolation between the two anchor points, CITED2 and VP16, increases model sensitivity. 1. [Line vs. Corner](https://drive.google.com/file/d/1foAbilR5Z3EXXWktCnCXBeONVcGruyyP/view?usp=sharing)

|  | Number of Predictions | Number of predictions on the gold standard list | Proportion of predictions on the gold standard list | p value (permutation) |
| --- | --- | --- | --- | --- |
| Original predictor | 144 | 26 | 0.181 | < 1e-4 |
| Original Predictor plus Region D (corner) | 312 | 37 | 0.119 | < 1e-4 |

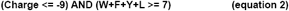

Region D contains 285 predictions, including 168 new predictions and 11 additional activation domains on the gold standard list (**Table 2**). Next, we lowered the W+F+Y+L boundary (**Figure 4A, Region E**) and lowered the acidic boundary (**Figure 4A, Region F**). When we consider only acidic activation domains, the projected tiles look very similar to those of the full gold standard list (**Figure S7**). Regions D and F contain the most new predictions.

### Sequence grammar

We attempted to improve our predictor by adding sequence grammar, which we define as the arrangement of amino acids. Examples of strict grammar include short linear sequence motifs (SLiMs), where amino acids must have a defined spacing or arrangement, e.g. ΦxxΦΦ, where Φ is a bulky hydrophobic residue (Dyson and Wright, 2016; Warfield et al., 2014). Examples of weak grammar include cases where acidic residues make larger contributions to activity when they are close to hydrophobic residues, or [DE][WFY] “mini motifs” that contribute to activity (Erijman et al., 2020; Ravarani et al., 2018; Staller et al., 2018). We saw that tiles with long runs of acidic residues were less likely to be activation domains (**Figure S8**). Otherwise, we could not detect statistically significant grammar signals, an outcome similar to those of other studies that have argued there is little grammar in activation domains (DelRosso et al., 2022; Erijman et al., 2020; Sanborn et al., 2021). The grammar that does exist is highly degenerate and flexible, making it hard to detect with our small sample size. Ultimately, we did not add grammar to the mechanistic predictor.

### Combining the mechanistic predictor with neural networks improves performance

We found that combining our mechanistic predictor with convolutional neural network predictors trained on yeast activation domains improved predictive power beyond the performance of either alone. Intersecting our predictor (n=144) and PADDLE (n=604) increased sensitivity (**Figure 5A**). For the 89 activation domains predicted by both models, 25 (28.1%) were on the gold standard list and 44 (49.4%) were on the Soto list (**Table 4, Table S7**). In addition, 88 had been tested in our activation domain assay and 45 (51.1%) had activity (Staller et al., 2022). This result implies each predictor brings orthogonal information. The 60 predictions removed by this intersection have many runs of acidic residues, consistent with the grammar analysis above.

**Table 4:**
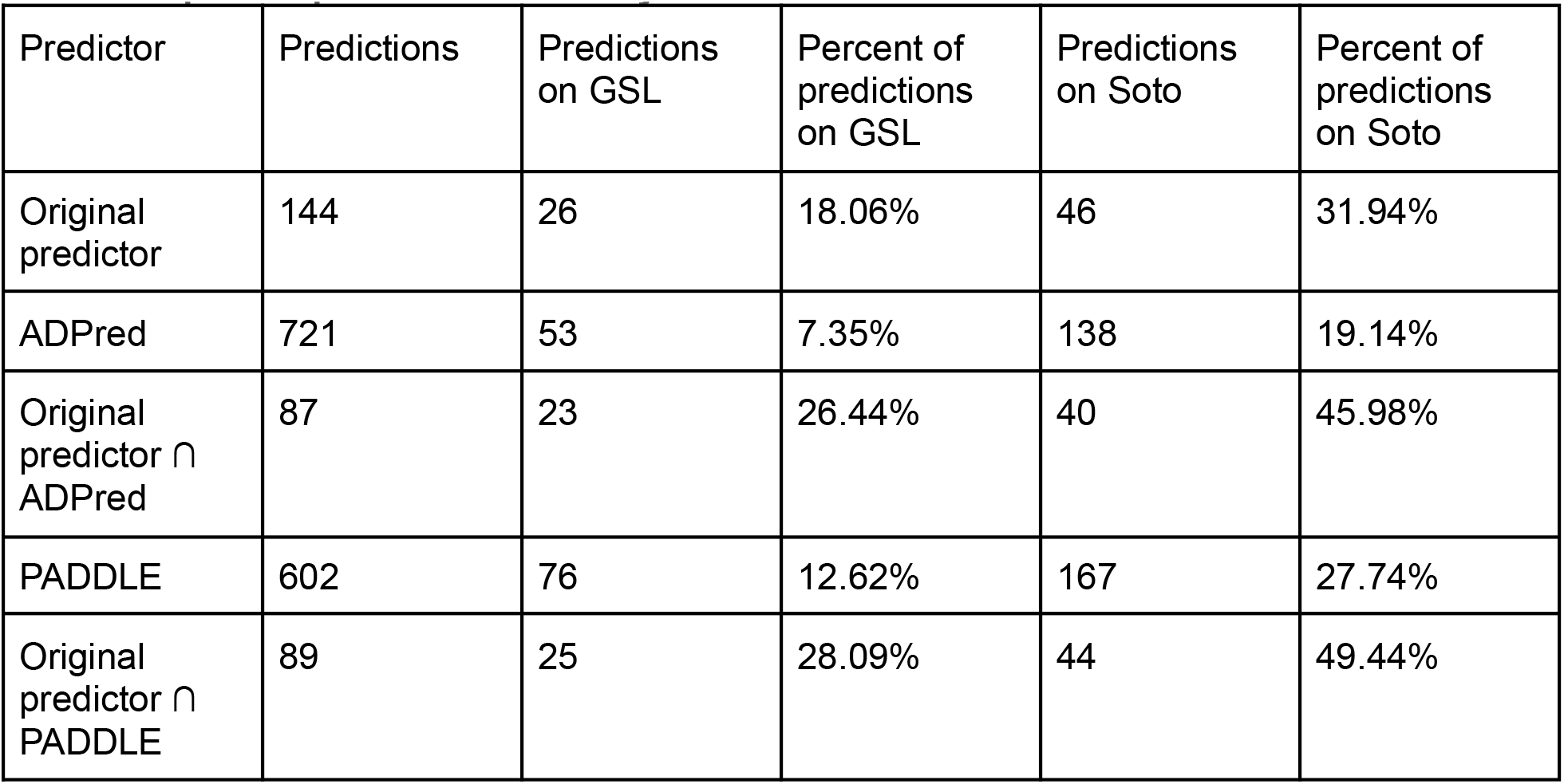
Intersection of the mechanistic predictor and convolutional neural network models improves prediction accuracy.

| Predictor | Predictions | Predictions on GSL | Percent of predictions on GSL | Predictions on Soto | Percent of predictions on Soto |
| --- | --- | --- | --- | --- | --- |
| Original predictor | 144 | 26 | 18.06% | 46 | 31.94% |
| ADPred | 721 | 53 | 7.35% | 138 | 19.14% |
| Original predictor ∩ ADPred | 87 | 23 | 26.44% | 40 | 45.98% |
| PADDLE | 602 | 76 | 12.62% | 167 | 27.74% |
| Original predictor ∩ PADDLE | 89 | 25 | 28.09% | 44 | 49.44% |

**Figure 5:**
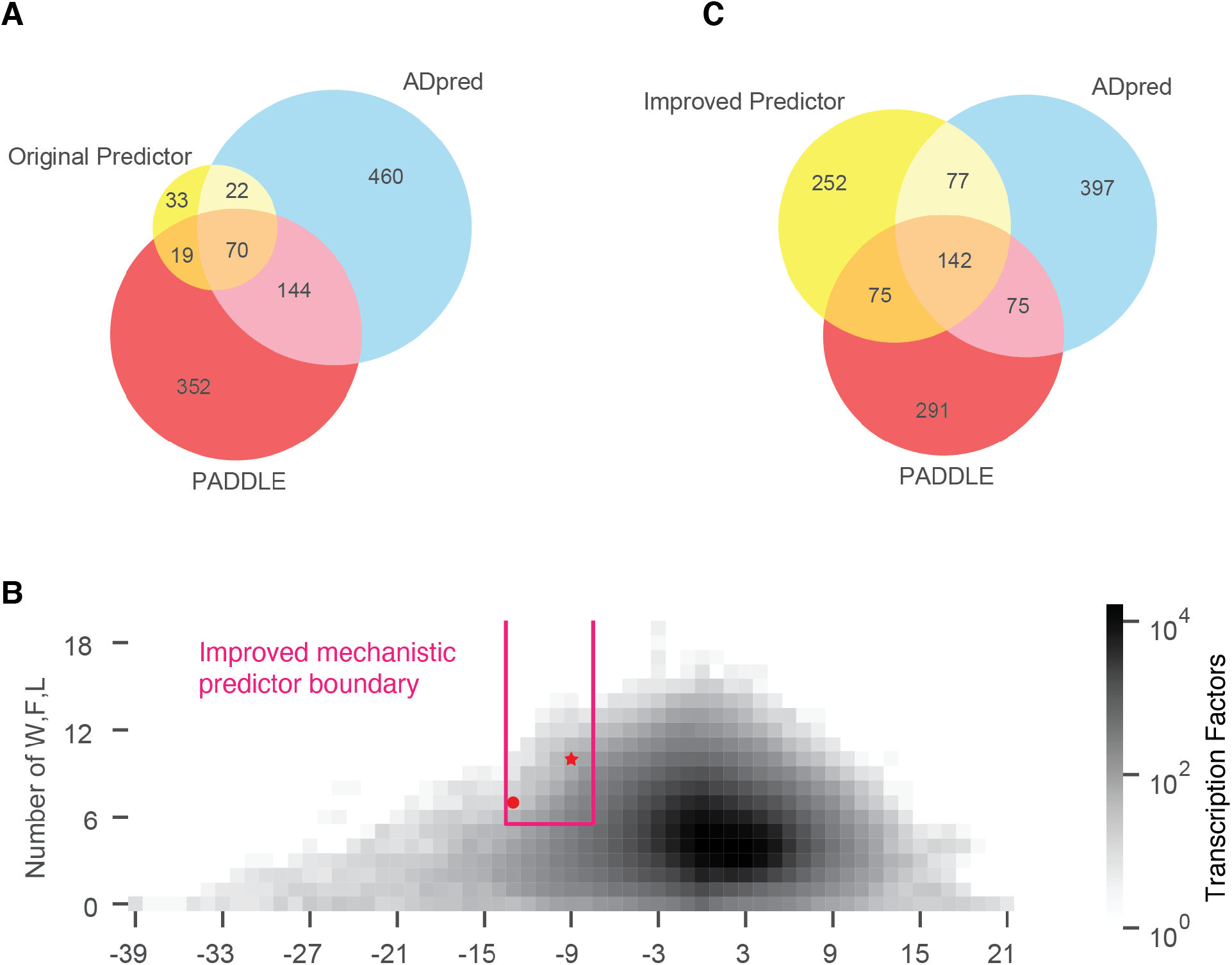
Intersections of the mechanistic predictors with the published neural network models from yeast. A) The overlap between the original mechanistic predictor, ADpred, and PADDLE predictions. B) The region of the improved predictor. C) The overlap between the improved mechanistic predictor, ADpred, and PADDLE predictions.

We found similar predictive improvement when we intersected our mechanistic predictor with ADpred. ADpred made 721 predictions on human transcription factors. Twenty-seven of these are on the gold standard list, and 45 overlap with the Soto list (**Table 4**). Intersecting the ADpred predictions with our predictor led to 87 overlaps: 23 (26.4%) with the gold standard list and 40 (46.0%) with the Soto list (**Table 4**). We had tested 86 of these regions in our experiments, and 40 (46.5%) had detectable activity in our assay (Staller et al., 2022). The intersection once again was more accurate than either model alone. ADpred and PADDLE scores are correlated (**Figure 5A**). We conclude that combining the convolutional neural networks and our mechanistic predictor yields the most accurate predictions.

Notably, there are five true activation domains found by our original predictor that are thrown out by PADDLE and ADpred. These activation domains from FOS, TIGD7, ZN513, TIGDF, and ZN777 contain many leucines (>10%), which is interesting because leucines make larger contributions to activity in human activation domains than in yeast activation domains (Staller et al., 2022). Indeed, based on our 15% threshold, FOS and ZN513 qualify as leucine-rich regions. We hypothesize that these leucine-rich activation domains are a metazoan innovation that bind to activation domain binding domains not present in yeast, such as the TAZ1 and TAZ2 domains of CBP/p300.

### An improved mechanistic predictor

We report an improved mechanistic acidic activation domain predictor (**Figure 5B**). We selected a trapezoidal region (equation 3), used seven amino acids (W,F,L,D,E,R,K) and expanded the charge and hydrophobic thresholds by one:

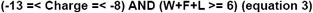

The trapezoid better emphasizes the balance between hydrophobic and acidic residues (**Figure 5B**). This region predicted 546 activation domains. Of these 546 predictions, 47 are on the gold standard list and 51 regions have high activity in our assays (**Table 5**). Moreover, 104/546 of our predictions are on the Soto list. The improved predictor identified 47/105 acidic activation domains on the gold standard list (44.8% sensitivity) and 104/290 acidic activation domains on the Solo list (35.9% sensitivity).This improved predictor makes 406 new predictions on 342 transcription factors. These transcription factors hail from a diverse set of families including many nuclear hormone receptors, Sox, Klf and Zinc finger transcription factors.

**Table 5:** Comparison of Soto and Gold Standard Lists. Source: [Table 1 Notebook](https://drive.google.com/file/d/1LOsCeiOj4k6CHT9QNkyBIhRja4Px1cae/view?usp=sharing)

|  | Number of predictions | Number of predictions overlapping the gold standard list | Proportion overlapping the gold standard list | Number overlapping the Soto list | Proportion overlapping the Soto list |
| --- | --- | --- | --- | --- | --- |
| Original predictor | 144 | 26 | 0.181 | 46 | 0.3194 |
| Improved predictor | 546 | 47 | 0.086 | 104 | 0.19 |

Intersecting this revised predictor with the convolutional neural networks yielded 139 high confidence predictions (**Table 6, Table S8**). We anticipate that testing this new set of predictions will identify new activation domains.

**Table 6:** Intersection of the improved mechanistic predictor and convolutional neural network models improves prediction accuracy.

| Predictor | Predictions | Predictions on GSL | Percent of predictions on GSL | Predictions on Soto | Percent of predictions on Soto |
| --- | --- | --- | --- | --- | --- |
| Improved predictor | 546 | 47 | 8.61 | 104 | 19.05 |
| ADPred | 721 | 53 | 7.35 | 138 | 19.14 |
| Improved predictor ∩ ADPred | 216 | 35 | 16.20 | 74 | 34.26 |
| PADDLE | 602 | 76 | 12.62 | 167 | 27.74 |
| Improved predictor ∩ PADDLE | 217 | 44 | 20.28 | 86 | 39.63 |

### The predictors identify one subclass of acidic activation domains

Both our original mechanistic predictor and the improved predictor do not identify all of the acidic activation domains on the gold standard and Soto lists.The true positive rate of the original (26/105) and improved predictors (47/105) are, strictly speaking, not amazing. Similarly, neither ADpred (53/105) nor PADDLE (76/105) can detect all these acidic activation domains. We have tuned the mechanistic predictors to have a low false positive rate at the expense of a high false negative rate. While there are multiple interpretations of this result, we favor the interpretation that there are multiple subclasses of acidic activation domains and that the existing predictors can find the one subclass that is well described by the acidic exposure model. These models miss activation domains where activity is regulated by modifying the net charge with post translational modifications. For example, p53 AD1 (net charge = -6, WFYL = 8) has the sites that increase activity when phosphorylated (S15, T18, S20, net charge = -12, WFYL = 8) (Raj and Attardi, 2017). These residues are interspersed with the aromatic and leucine residues consistent with the acidic exposure model. In this case, the resting sequence falls outside the activation domain predictor boundary (equation 3), but the activated, phosphorylated state crosses it over into the active domain. Understanding how phosphorylation controls activation domain function is an exciting area of future inquiry.

## Discussion

Accurate computational models for predicting activation domains from protein sequence will advance basic science and precision medicine. Computationally annotating activation domains would allow studies of how paralogous transcription factors diversify after duplication and enable evolutionary comparisons of domain shuffling. Comprehensive lists of transcription factors with activation domains could improve gene regulatory networks by adding signs to the connections inferred from genome binding data (Hummel et al., 2022) or by distinguishing direct and indirect connections inferred from genetic perturbations. Predicting activation domains is a key step towards building models that predict how mutations in activation domains modulate activity, which, in the long term, could classify patient mutations in activation domains as benign or pathogenic (Richards et al., 2015; Starita et al., 2017). These classifications could group patients for the development of targeted therapies or prioritize variants for base-editing gene therapies.

Our mechanistic predictor is valuable because it is simple and interpretable. Its accuracy comes from the acidic exposure model, which describes a subclass of acidic activation domains that balance acidic residues with key hydrophobic residues. The predictor’s success further supports the acidic exposure model. Our acidic exposure model is related to the stickers and spacers model for how some specialized IDRs mediate condensate formation (Martin et al., 2020), albeit with a more active role for the spacers. So far, we can predict only acidic activation domains. Analogous predictors of P-rich, Q-rich, or S-rich activation domains do not work (**Figure S4, S5, S6**).

Why are so many activation domains negatively charged? What is the mechanistic advantage of acidity? This question has been repeated many times since it was posed by Paul Sigler (Sigler, 1988). In principle, exposure of hydrophobic residues could be achieved by positively charged residues, but, in practice, positively charged residues inhibit activation domain function (Arnold et al., 2018; Erijman et al., 2020; Ravarani et al., 2018). Many coactivators have positively charged surfaces, and long-range, low-affinity fast electrostatic interactions have been documented (Almlof et al., 1997; Hermann et al., 2001). These electrostatic interactions can be important for making activation domain coactivator interactions diffusion-limited in “fly-fishing” models of activation domain coactivator interactions (Kim et al., 2018; Kim and Chung, 2020). We believe there are advantages to acidic activationd domains and disadvantages to basic activation domains. For the advantages, first, acidic residues electrostatically repel the DNA, allowing the activation domain to stick out and catch coactivators. Second, acidic activation domains can have affinity electrostatic intramolecular interactions with basic DNA binding domains, which can increase the specificity of DNA binding via competitive inhibition (He et al., 2019; Krois et al., 2018; Stott et al., 2014; Wang et al., 2021). Third, acidity makes it possible to post-translationally regulate activation domain activity with phosphorylation (Conti et al., 2023). We see two potential disadvantages to basic activation domains: first, nonspecific, electrostatic binding to DNA that could compete with coactivator binding or inhibit nuclear search. Second, cation-π interactions between basic residues and aromatic residues (e.g. arginine-tyrosine interactions (Wang et al., 2018) could drive collapse (or condensate formation) and make positively charged residues less effective at keeping some aromatic residues exposed to solvent. There is also some evidence that some repression domains are positively charged. Together, these observations explain why so many activation domains are acidic.

Activation domains display very flexible sequence grammar. If activation domains had strict sequence grammar requirements for function, we would have seen these signatures in the evolutionary record, mutagenesis, or in tiling experiments. An early grammar model, the 9aaTAD model, can identify known activation domains, but in high-throughput screens of random peptides or yeast transcription factors, it does not detect more often than expected by chance (Erijman et al., 2020; Piskacek et al., 2007; Sanborn et al., 2021). Instead, we see evidence for very flexible grammar or no grammer. The evidence for no grammar is that random peptides can have activation domain activity and that shuffling activation domain sequence can preserve or even sometimes increase activity (Arnold et al., 2018; Erijman et al., 2020; Ma and Ptashne, 1987; Ravarani et al., 2018; Sanborn et al., 2021; Staller et al., 2018). The high accuracy of our grammar-less composition-based predictor supports both a no-grammar model and a flexible-grammar model. The evidence against no-grammar models is that shuffling activation domain sequence can both increase and decrease activity (Sanborn et al., 2021; Staller et al., 2018). Loss of activity is more common when shuffling disrupts an alpha helix (Sanborn et al., 2021; Staller et al., 2022). In these shuffle mutants, the arrangement of amino acids, i.e. the grammar, is modulating activity, ruling out a strict no-grammar model. We can rule out a strict-grammar model and we can rule out a no-grammar model, so we are left with a very-flexible-grammar model.

How do we square a very-flexible-grammar with the documented role of short linear motifs? The dominant model for activation domains is that they are anchored by a hydrophobic short linear motif embedded in a permissive context. At this time, the features of the context are more clearly defined than the motifs. The context is acidic residues and an intrinsically disordered region. In some cases, the motifs are clearly present, conserved, and contribute to activation domain activity (Dyson and Wright, 2016). A motif in an amphipathic alpha helix is a very effective way to coherently display several hydrophobic residues to a coactivator (Giniger and Ptashne, 1987). Amphipathic alpha helices are a good solution for building an activation domain (Dyson and Wright, 2016). They are not the *only* solution. Motifs are uncommon and rarely generalize beyond a few TFs. Surveys of random peptides, yeast transcription factors, and human transcription factors found enrichment of only [DE][WFY] ‘mini motifs’ (Arnold et al., 2018; DelRosso et al., 2022; Erijman et al., 2020; Ravarani et al., 2018; Sanborn et al., 2021). We argue that the critical distinction is that a motif or an amphipathic helix is not the only way for a cluster of hydrophobic residues to interact with a coactivator–many arrangements are functional. A growing number of fuzzy interactions have been documented, but they are likely underreported because of investigator bias and a higher burden of proof (Brzovic et al., 2011; Risør et al., 2021; Tuttle et al., 2018; Warfield et al., 2014). Fuzzy binding is consistent with a highly-flexible-grammar.

It is not clear at what point the motif ends and the context begins. In our mutagenesis of VP16 and CITED2, we found that virtually every hydrophobic residue contributed to activity, blurring the distinction between motifs and context. Adding aromatic residues near a motif – in essence extending the motif – increases activation domain activity (Staller et al., 2018; Warfield et al., 2014). Based on mutagenesis of Abf1, one group has argued that motif quality and context quality both contribute to function and that each can compensate for the other (Langstein-Skora et al., 2022). Sequences that contain many functional elements will be composition driven and grammar-independent (e.g. DWDWDWDWDWDWDWDWDWDW (Ravarani et al., 2018)). Grammar and motifs will be important on the margin for sequences that barely have the right composition to be functional but can function when the residues are appropriately arranged into a motif. Regions with fewer acidic and hydrophobic residues will likely be more reliant on grammar. Critically, even in a highly-flexible-grammar regime, not all arrangements of residues will be active. Real sequences are likely to be on this margin because neutral drift is likely pulling strong activators down to the minimum functional level. Marginally active activation domains would be easier to regulate by post-translational modifications. Weak or regulated activation domains allow more precise combinatorial control of gene expression. We speculate that there is no boundary between motifs and context.

## Conclusion

We conclude that the human proteome contains a class of strong acidic activation domains that requires balance between W,F,L and acidic residues. This balance accurately predicts known activation domains, many of which are well described by our acidic exposure model. Our work implies there are other classes of acidic activation domains that are not predicted by our model and which likely bind to other coactivators. Forthcoming, comprehensive maps of activation domains (DelRosso et al., 2022) will create the opportunity to test and improve the mechanistic predictors. Our analyses emphasize the need to characterize the sequence features that control activity of Q-rich, S-rich, and P-rich activation domains.

## Acknowledgments

The authors thank Jordan Stefani, Zeba Wunderlich, Alex Holehouse, Vinson Fan, and Darren Kahan for comments on the manuscript. Thanks to Shiyi Yang and Angelica Lam for help for setting up and running ADpred.

## Funding

S.K. is an undergraduate at UC Berkeley supported by the UC Berkeley STEM Excellence through Equity & Diversity Scholars Honors Program. This work was supported by the Burroughs Wellcome Fund Postdoctoral Enrichment Program and NSF grant 2112057.

## Author Contributions

S.K. Conceptualization, Software, Formal analysis, Writing. M.V.S. Conceptualization, Methodology, Writing.

## Conflict of Interest

The authors declare no competing interests.

## Materials and Methods

### Data sources

The gold standard list consists of activation domains from two sources. The first source is transcription factors with activation domains that were annotated in Uniprot, which are in the “data/UniprotActivationDomains_HighqualitySet.csv” file. The second source is activation domains manually curated from the literature, which are in “/data/ActivationDomainsHuman.csv”. In “/notebooks/Building the GSL.ipynb,” we combined activation domains from both sources into one list, merging entries with the same uniprotID with overlapping start and end positions. In cases where the annotated boundaries differed, we used the longest region.

For the Tested Regions in **Figure 1** and **Figure 4**, we used the activity data for fragments that we previously published (Staller et al., 2022). There were 443 entries, but not all were recovered in the assay. In addition, there were multiple cases where the predicted activation domain region was different from the Uniprot region, and we experimentally tested both. Here we combined overlapping regions, yielding a total of 356 regions, which include positive controls, negative controls, and predicted activation domains.

The sequences of all 1608 human transcription factors were downloaded from Uniprot (Lambert et al., 2018). We downloaded the sequences of the full human proteome from Uniprot. For analysis, we used the protfasta (Holehouse, 2021), localcider (Holehouse et al., 2017), metapredict2 (Emenecker et al., 2022), pandas, math, itertools, matplotlib, seaborn, numpy, and scipy python packages.

### Tiling

Each transcription factor sequence is computationally chopped into 39AA regions spaced every 1AA (e.g. 1-39, 2-40, 3-41 etc.), yielding 881K tiles. The full proteome and gold standard list were similarly decomposed into tiles. The composition of each tile was computed by counting amino acids. Net charge was calculated as R+K-D-E. The original predictor counted W+F+Y+L residues. Other variants of the predictor counted other subsets of residues. For the composition histograms in **Figure 2**, we used the tiles. To identify enrichment of specific amino acids in activation domains, we performed a student’s t-test between the two population means. We report all enrichments that were significant at p <0.01 after a Bonferroni correction for multiple hypothesis testing.

### Varying the predictor

The main predictor function is make_predictions () in “notebooks/AD_predictor_tools.py.” The make_predictions () function has parameters that can be used to vary the predictor. We varied the predictor in one way at a time. We adjusted the parameter UpperCorner_slope1 to change the slope of the boundary line originating from the upper corner, CITED2. To create region D in **Figure 4A**, we set UpperCorner_slope1 equal to infinity to draw a vertical line to test the predictor with a vertical boundary line instead of a diagonal line. To vary the acidity threshold, we adjusted the parameter upper_corner_c from -13 to 0 to vary the charge of the boundary. To vary the hydrophobic residue count, we adjusted the parameter lower_corner_h from 0 to 15 to vary the count of hydrophobic and aromatic AAs. To vary the composition of AAs used on the y-axis we substituted each pair and triad of AAs for the composition parameter.

We identified all the tiles that satisfied equation 1 and then aggregated the overlapping tiles to predict activation domains. For longer predictions, there can be tiles in the middle that do not individually satisfy equation 1. As a result, when we project the tiles from our tested predictions (**Figure 4B**), some of the tiles are outside the prediction region. We initially allowed overlaps of ≥1 AA, but in practice, the minimum observed overlap was 26 residues. For each version of the predictor, we recalculated the properties, changed the inequalities, found new sets of tiles, aggregated the tiles into predictions, and compared the predictions to the gold standard list.

We used the function compare_to_random () in “notebooks/AD_predictor_tools.py” to compare the overlap with the gold standard list of predictions vs random sequences. To create a random set of sequences, we counted the number of unique uniprotIDs in the predictions and randomly sampled the same number of transcription factors from all human transcription factors. Then, for each unique uniprot ID in the predictions, we recorded the length distribution and number of predictions. We iterated through unique uniprotIDs and randomly selected parts of each sampled transcription factor so that the selected parts had the same number and length distribution as predictions with one uniprot ID. Finally, we recorded the number of times a randomly sampled transcription factor region had any overlap with a gold standard list entry, using start, end, and uniprotID to compare. Here we used 1 AA overlaps to increase the stringency of the permutation test. We compared this number to overlaps of predictions with the gold standard list. We repeated this random sampling for 10,000 permutations. We never randomly sampled as many overlaps as the predictor.

### Comparing predictions to convolutional neural networks

The two neural networks to which we compared the predictor were ADPred and PADDLE (Erijman et al., 2020; Sanborn et al., 2021). We installed ADPred from https://github.com/FredHutch/adpred and then ran it on all Lambert transcription factors. Using the criteria detailed at https://adpred.fredhutch.org/, we considered sequences with at least 10 consecutive positions with a score of at least 0.8 to be a predicted activation domain. We downloaded the activation domains predicted on human transcription factors by PADDLE at https://cdn.elifesciences.org/articles/68068/elife-68068-fig3-data1-v3.xlsx. We used predictions of both medium and high strength.

We used the compare_two_predictors () function in “notebooks/AD_comparison_tools.py” to compare our predictor to the neural net predictors. This function first individually compares the overlap of our predictions and a neural net’s predictions with a list of known activation domains (either the gold standard or the Soto et al. list). Then, it compares our predictions that overlap with a prediction made by a neural network to the list of known activation domains.

The predicted activation domain regions are deposited at https://zenodo.org/badge/latestdoi/548126430

The analysis code was deposited at https://zenodo.org/badge/latestdoi/548126430

### Supplementary Data Tables

Table S1: The gold standard list of activation domains.

Table S2: Removing each of the original eight AA from the predictor did not improve performance.

Table S3: Varying the length of the tiles did not improve performance. Table S4: Using single amino acids as the y-axis of the predictor.

Table S5: Using pairs of amino acids as the y-axis of the predictor. The top 20 pairs are reported.

Table S6: Using triplets of amino acids as the y-axis of the predictor. The top 20 triplets are reported.

Table S7: Activation domains predicted by both our original predictor and the yeast neural networks.

Table S8: The coordinates and sequences of the activation domains predicted by the revised predictor.

## Notes

### Competing Interest Statement

The authors have declared no competing interest.

