## SupplementalFigures for "The balance of acidic and hydrophobic residues predicts acidic transcriptional activation domains from protein sequence"

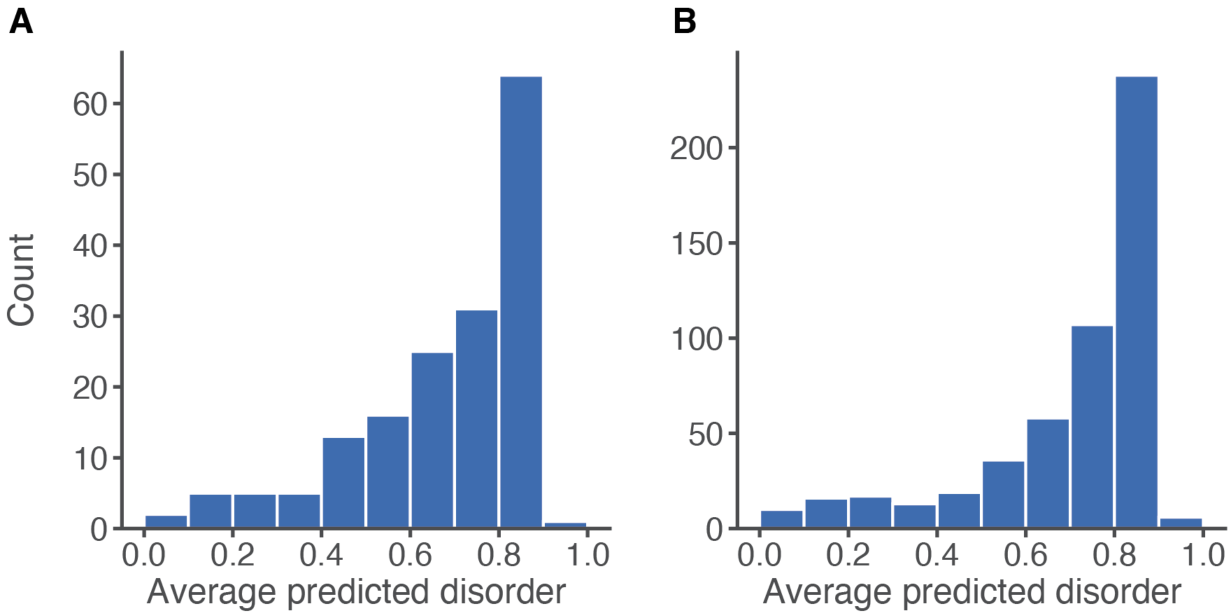


**Figure S1:** The majority of activation domains from the gold standard list (A) and the Soto list (B) are predicted to be intrinsically disordered with Metapredict2 (Emenecker et al., 2022). For each full length activation domain, we computed the Metapredict2 score for each residue and then averaged it over all residues. Regions with a score above 0.5 are predicted to be disordered.

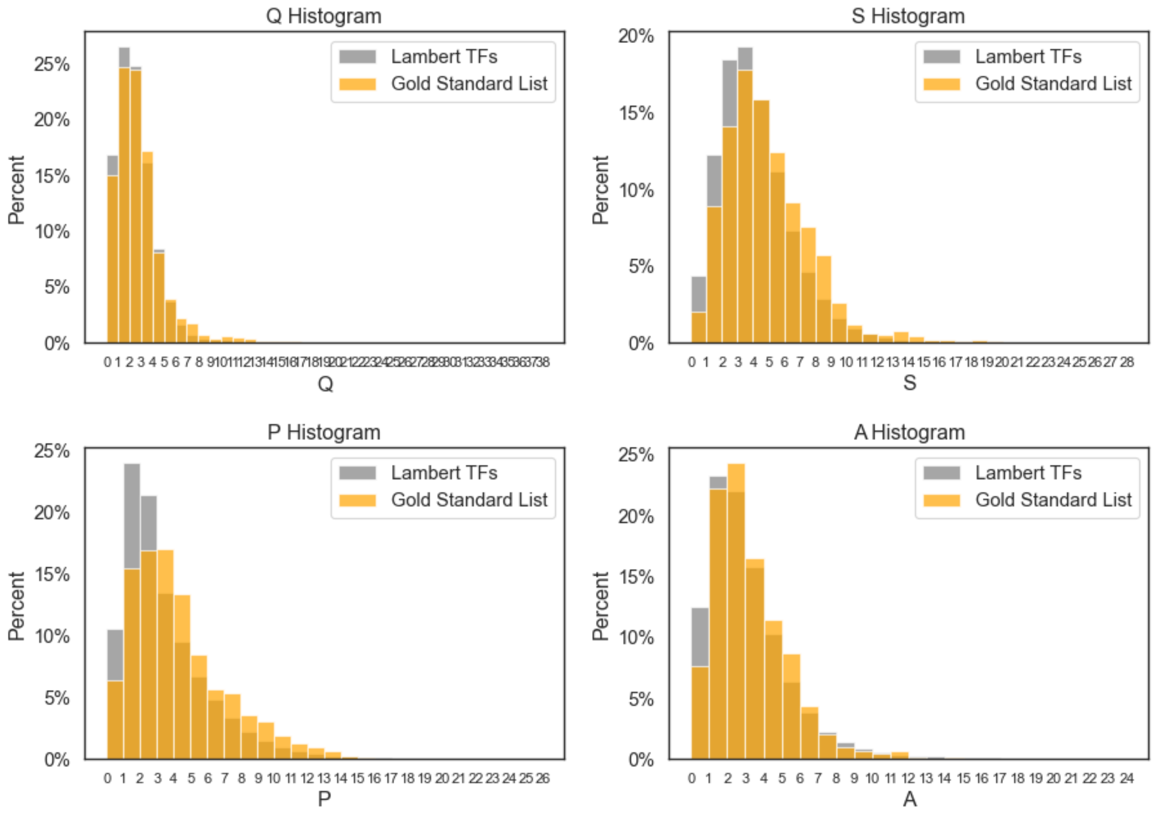


**Figure S2:** Histograms of additional sequence features of tiles of activation domains on the gold standard list.

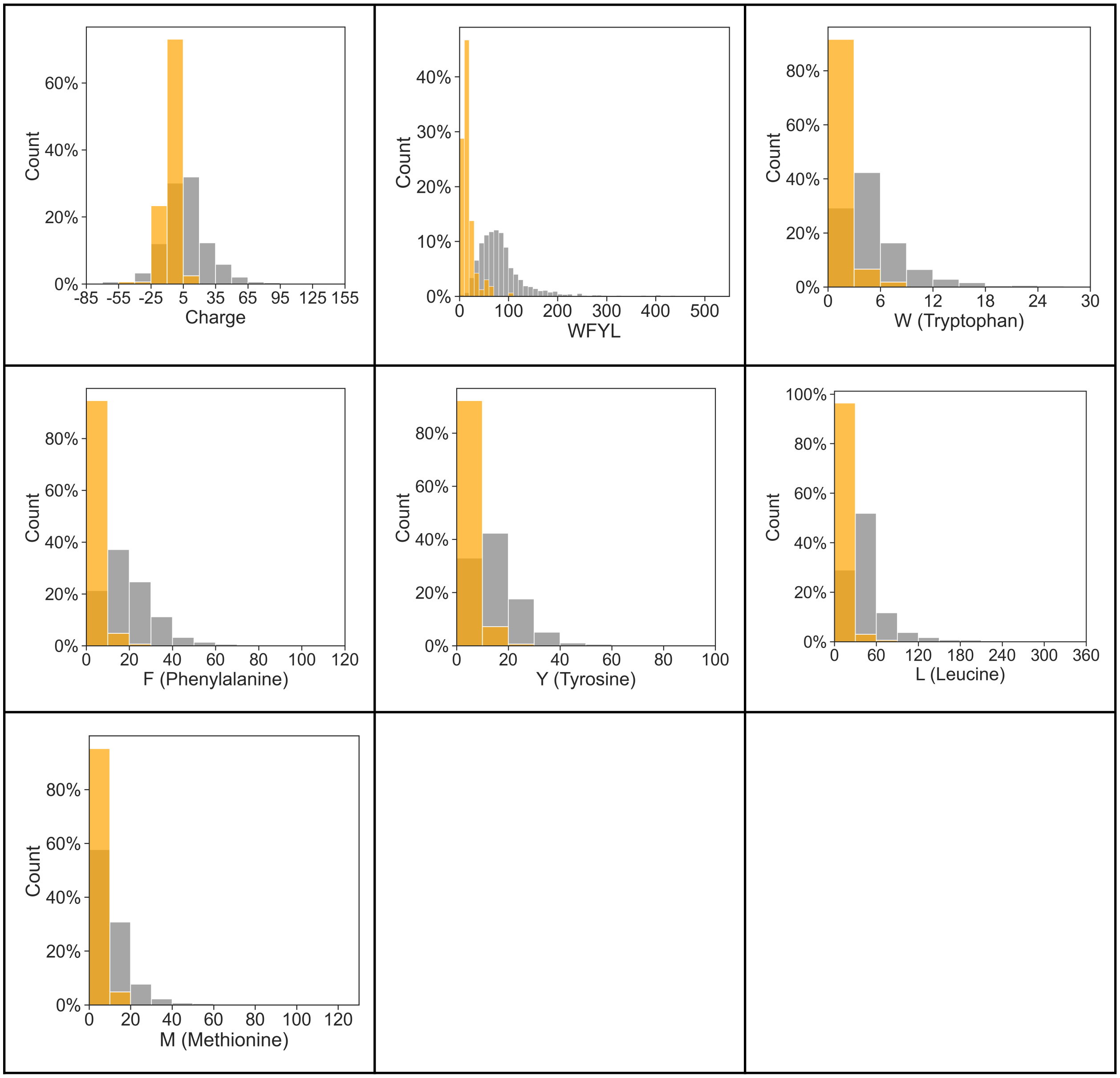


**Figure S3:** Histograms of amino acid frequencies in gold standard list activation domains before normalizing for length or computing tiles. Because all the activation domains had different lengths, these histograms are not particularly informative. This problem prompted us to decompose regions into 39-AA tiles in the main text and Figure S1. Gray, all TFs. Orange, gold standard list activation domains.

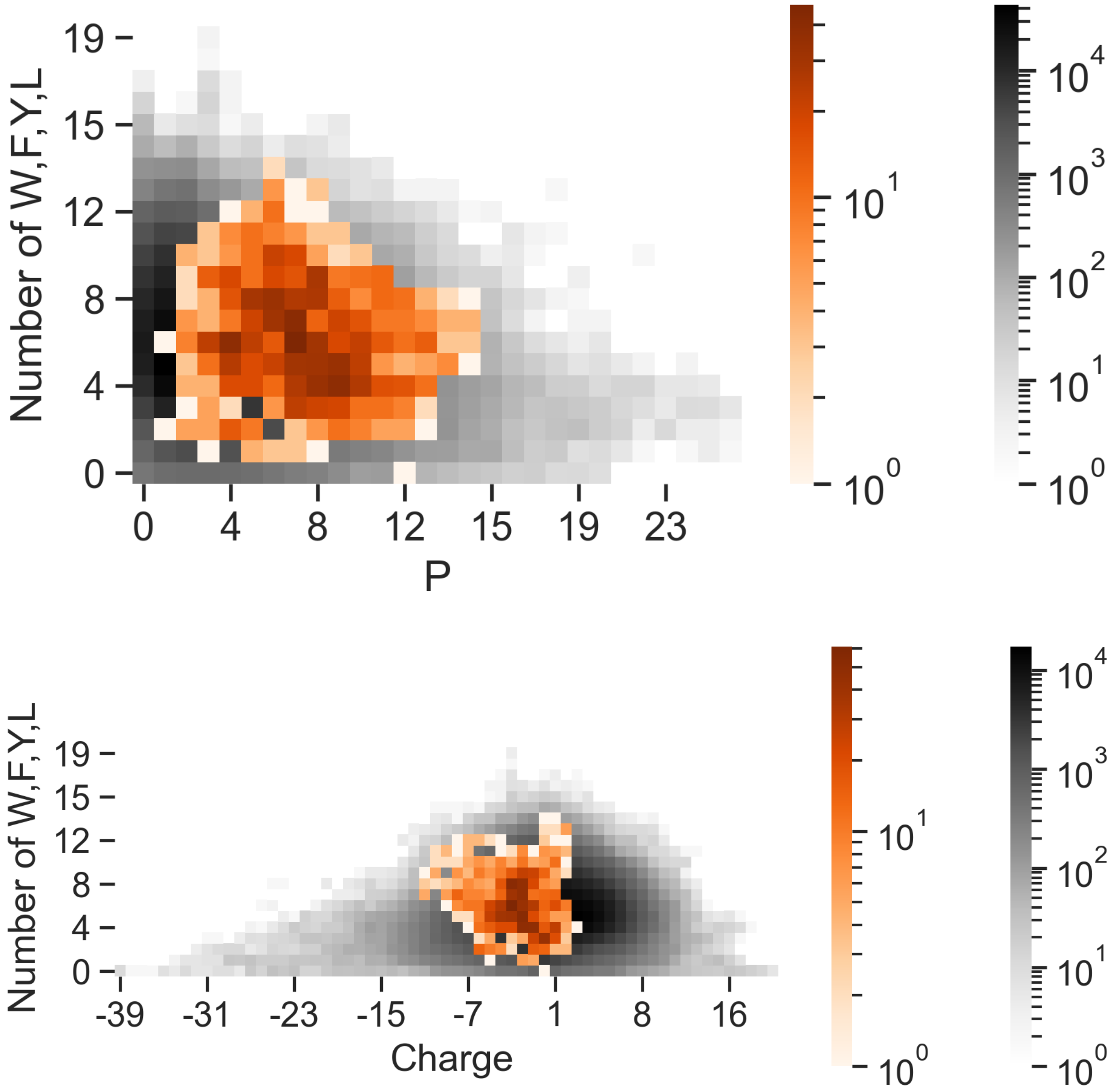


**Figure S4:** Proline-rich activation domains from the gold standard list are not among the most proline-rich regions of transcription factors. For all 881K tiles from TF, we counted P residues and WFYL residues. The combination of P and WFYL residues does not enrich P-rich activation domains. The tiles from P-rich activation domains often have negative charges, but are less negative than acidic activation domains.

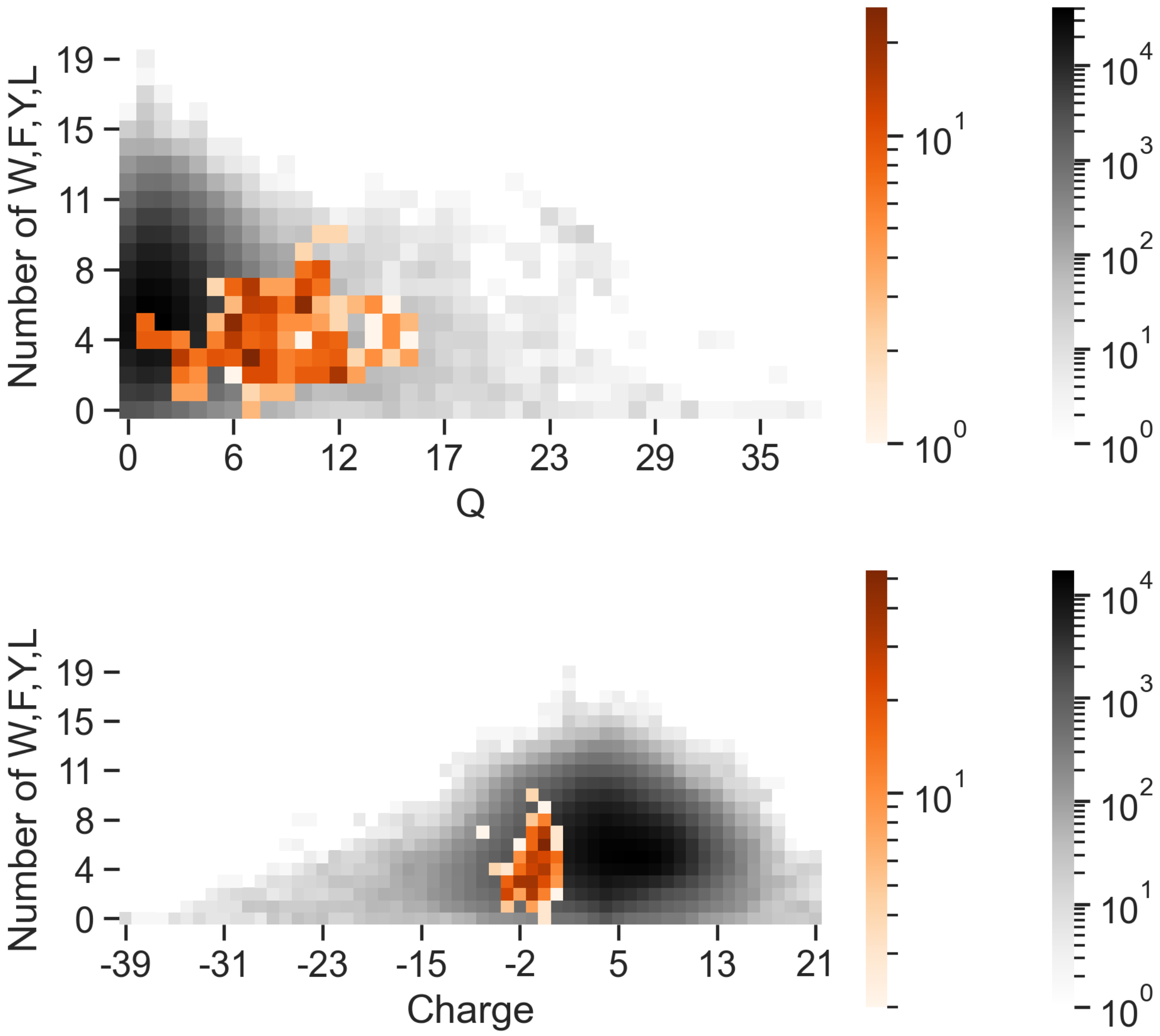


**Figure S5:** Glutamine-rich activation domains from the gold standard list are not among the most glutamine-rich regions of transcription factors. For all 881K tiles from TF, we counted glutamine (Q) residues and WFYL residues. The combination of Q and WFYL residues does not enrich Q-rich activation domains. Most tiles from Q-rich activation domains are near neutral or slightly acidic.

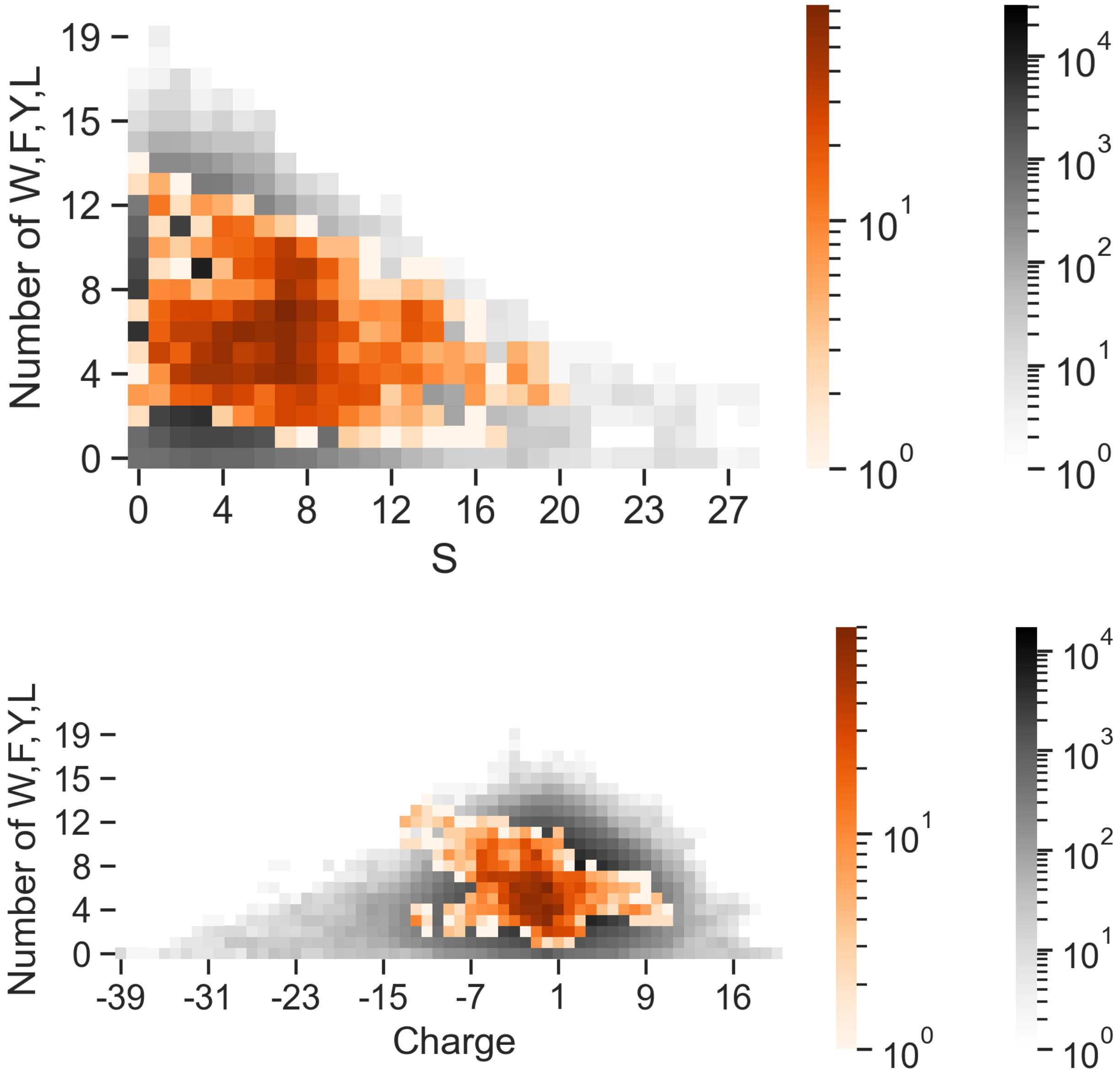


**Figures S6:** Serine-rich activation domains from the gold standard list are not among the most serine-rich regions of transcription factors. For all 881K tiles from TF, we counted glutamine (S) residues and WFYL residues. The combination of S and WFYL residues does not enrich S-rich activation domains. The tiles from S-rich activation domains show a wide range of net charge.

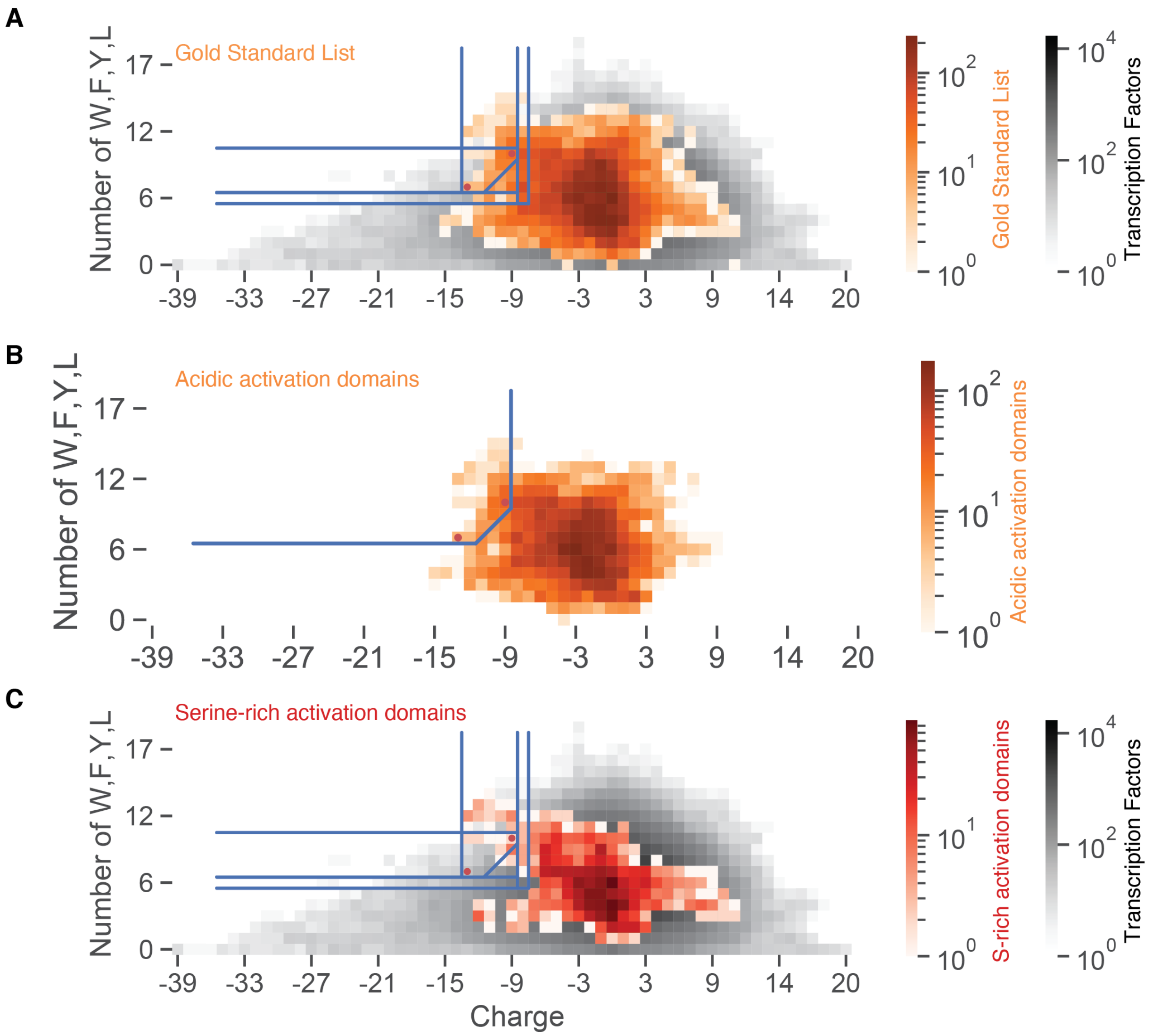


**Figure S7:** The distribution of sequence features in gold standard list acidic activation domains is the distribution of properties of all activation domains on the gold standard list.

A) Tiles from all activation domains from the gold standard list (n = 167).

B) Tiles from the acidic activation domains on the gold standard list (n = 105). There are fewer positively charged tiles.

C) Tiles from the S-rich activation domains on the gold standard list (n = 37). Unsurprisingly, the acidic activation domain predictor struggles to predict the majority of S-rich activation domains. S-rich activation domains are determined by a different mixture of sequence features.

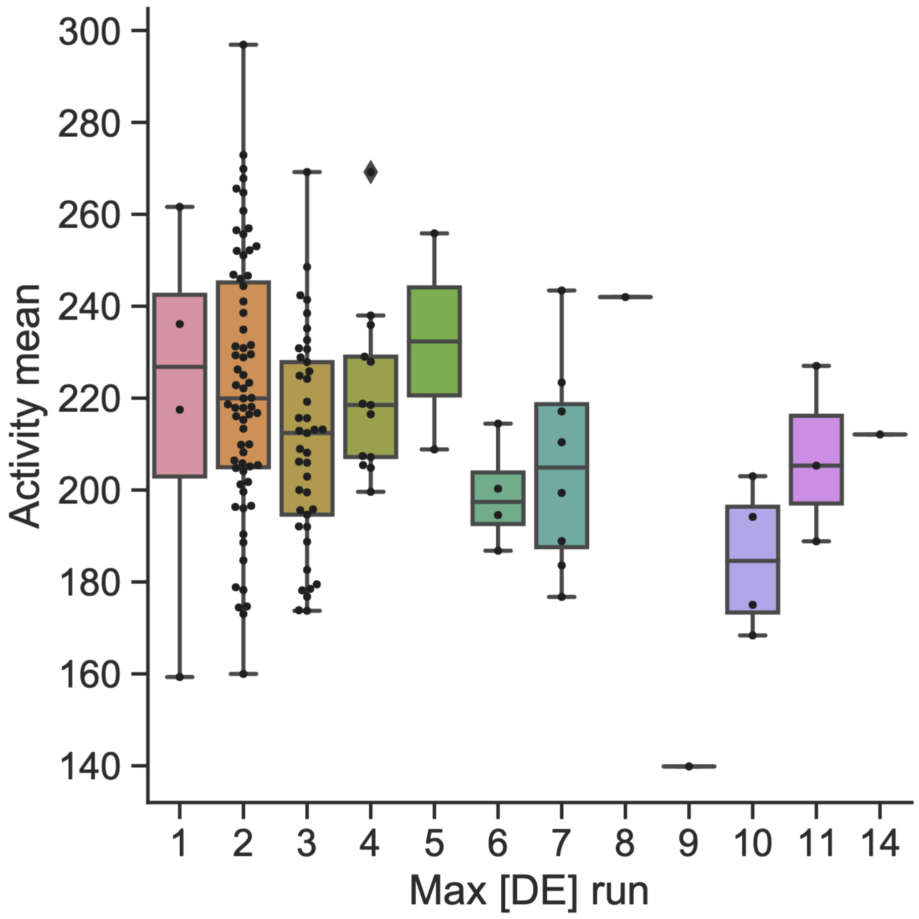


**Figure S8:** Long runs of acidic residues are depleted from strong activation domains. These are the tested predictions from the original predictor, replotting the data from Staller et al. 2022. For each activation domain, we counted the longest contiguous run of acidic residues (D or E) and plotted it against activity. In this experiment, the no activation domain control was normalized to 200. Strong activation domains had activity greater than 221. Boxplots show median and interquartile range, whiskers show full range of the data. Sequences with long runs of acidic residues tended to have lower activity, but the effect is weak.
